# Human DNA-PK activates a STING-independent DNA sensing pathway

**DOI:** 10.1101/587501

**Authors:** Katelyn Burleigh, Joanna H. Maltbaek, Stephanie Cambier, Richard Green, Michael Gale, Richard C. James, Daniel B. Stetson

## Abstract

Detection of intracellular DNA by the cGAS-STING pathway activates a type I interferon-mediated innate immune response that protects from virus infection and can be harnessed to promote anti-tumor immunity. Whether there are additional DNA sensing pathways, and how such pathways might function, remains controversial. We show here that humans – but not mice – have a second, potent, STING-independent DNA sensing pathway that is blocked by the E1A viral oncogene of human adenovirus 5. We identify human DNA-PK as the sensor of this pathway and demonstrate that DNA-PK kinase activity drives a robust and broad antiviral response. We discover that the heat shock protein HSPA8/HSC70 is a unique target of DNA-PK. Finally, we demonstrate that detection of foreign DNA and DNA damage trigger distinct modalities of DNA-PK activity. These findings reveal the existence, sensor, unique target, and viral antagonists of a STING-independent DNA sensing pathway (SIDSP) in human cells.

## Introduction

The cGAS-STING DNA sensing pathway has emerged as a key component of the innate immune response that is important for antiviral immunity (*1*), contributes to specific autoimmune diseases (*2*), and mediates important aspects of antitumor immunity (*3*). cGAS binds to double-stranded DNA and catalyzes the formation of cyclic GMP-AMP (*4, 5*), a diffusible cyclic dinucleotide that activates the endoplasmic adaptor protein STING (*6*). Activated STING then serves as a platform for the inducible recruitment of the TBK1 kinase, which phosphorylates and activates the transcription factor IRF3, leading to the induction of the type I interferon mediated antiviral response (*7*).

Nearly all studies on the cGAS-STING pathway involve the use of mice and mouse cells. Knockouts of cGAS (*Mb21d1*) and STING (*Tmem173*) have clearly demonstrated that both are essential for the transcriptional and cell biological responses to foreign intracellular DNA, and that they mediate the pathology of specific autoimmune diseases (*2, 8, 9*). However, numerous additional sensors of intracellular DNA have been proposed, all of which are thought to act upstream of STING (*10*). Whether STING-independent DNA sensing exists is currently unknown.

Here, we report the unexpected finding that the E1A oncogene of human adenovirus 5 blocks two distinct DNA sensing pathways in human cells: the well-known cGAS-STING pathway (*11*), and a second, STING-independent DNA sensing pathway (SIDSP). We identify the DNA damage response protein DNA-PK as the sensor of the SIDSP, along with the heat shock protein HSPA8 as a unique SIDSP target. We show that the SIDSP is potently activated in human and primate cells, but it is weak or absent from mouse cells. Our findings demonstrate that human cells have a second DNA sensing pathway, with implications for host defense, autoimmunity, and anti-tumor immunity.

## Results

### Human adenovirus 5 E1A blocks two DNA sensing pathways in human cells

We previously demonstrated that the viral oncogenes of the DNA tumor viruses are potent antagonists of the cGAS-STING DNA sensing pathway (*11*). We sought to define the mechanism of this antagonism, focusing on the E1A oncogene of human adenovirus 5, which is constitutively expressed in human HEK 293 cells and is responsible for their transformation. As shown previously (*11*), we found that HEK 293 cells mounted a robust type I IFN response to RIG-I ligand, but not to transfected calf thymus (CT) DNA, and that CRISPR-mediated disruption of E1A restored the DNA-activated IFN response (Fig. 1A). We monitored key early events in the antiviral signaling pathways that are activated by cGAS-STING and the RNA-activated RIG-I-MAVS pathways: the activation and autophosphorylation of the kinase TBK1, and the subsequent TBK1-dependent phosphorylation of the IRF3 transcription factor on several residues that leads to IRF3 dimerization, nuclear translocation, and transcription of the type I IFN genes (*7*). In E1A-expressing control HEK 293 cells, we found that CT DNA transfection resulted in detectable phosphorylation of TBK1 on serine 172, but phosphorylation of IRF3 on serine 396 was impaired compared to RIG-I ligand-activated IRF3 phosphorylation (Fig. 1B). Disruption of E1A restored DNA-activated IRF3 S396 phosphorylation (Fig. 1B). These data suggested that E1A blocks IRF3 activation at a step between TBK1 activation and IRF3 phosphorylation, leading us to evaluate IRF3 activation more thoroughly. To do this, we used an antibody to detect phosphorylation of IRF3 on serine 386, which is essential for IRF3 dimerization (*7*). To test for the specificity of the response, we used lentiCRISPR to generate clonal lines of STING- and TBK1-deficient HEK 293 cells, validating disruption of the genes by DNA sequencing and western blot (Fig. S1). We found that transfection of either DNA or the direct STING agonist cGAMP resulted in robust IRF3 S386 phosphorylation in E1A-expressing control HEK 293 cells, but IRF3 S396 phosphorylation was impaired (Fig. 1C). This IRF3 S386 phosphorylation after cGAMP treatment was absent from STING- and TBK1-deficient cells, confirming that cGAMP-induced IRF3 activation was both STING- and TBK1-dependent. Surprisingly, however, DNA-activated IRF3 S386 phosphorylation was intact in both STING-deficient and TBK1-deficient HEK 293 cells (Fig. 1C). Disruption of E1A restored both STING-dependent IRF3 S396 phosphorylation in response to cGAMP and STING-independent IRF3 S396 phosphorylation in response to DNA. These data demonstrate that E1A blocks the DNA-activated antiviral response at a late step, after initiation of IRF3 phosphorylation but before its completion. Moreover, E1A unexpectedly blocks two distinct antiviral responses in HEK 293 cells, one STING-dependent and one STING-independent.

**Figure 1:**
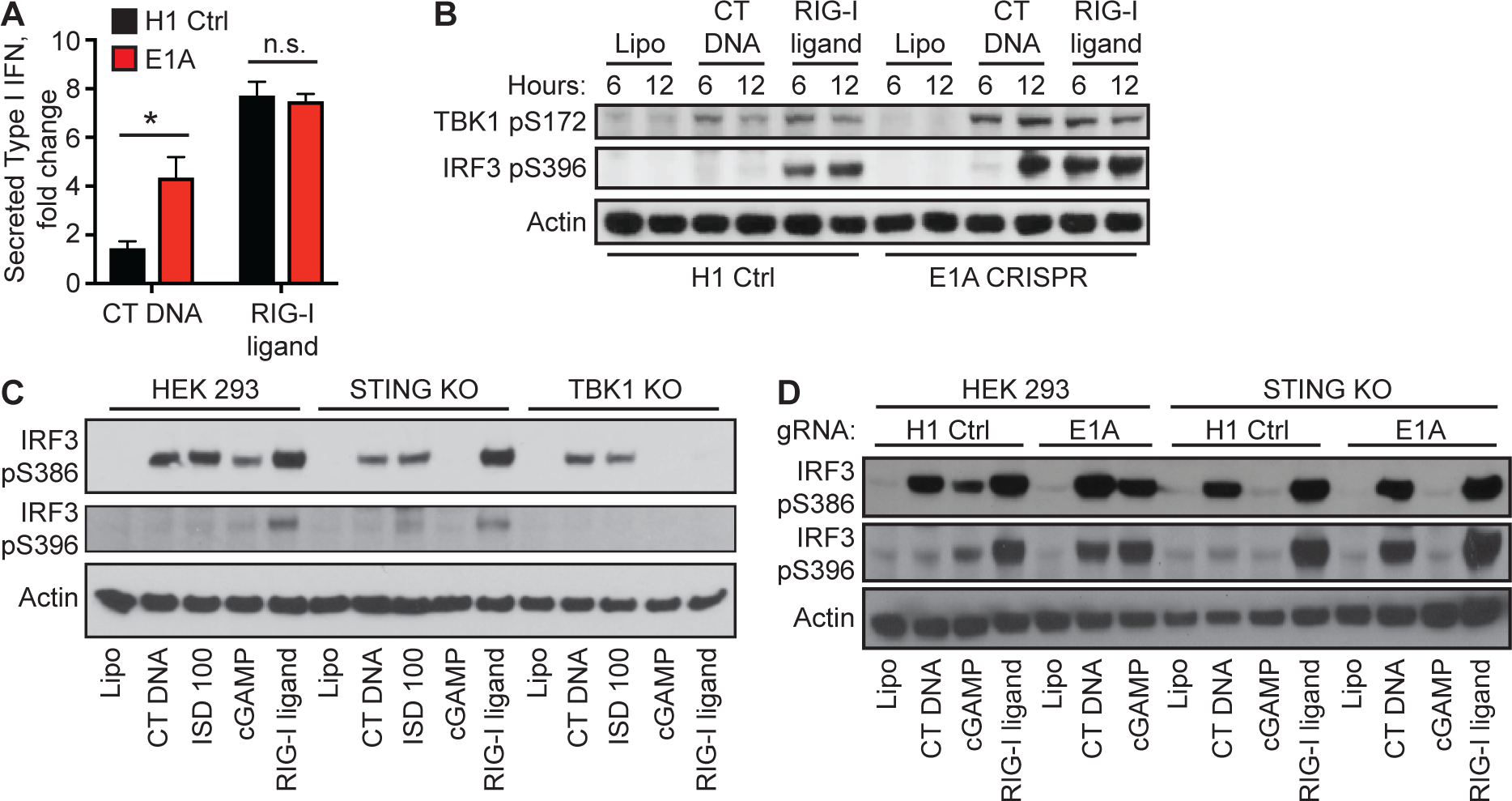
Human Adenovirus 5 E1A blocks two DNA sensing pathways. (A) HEK 293 cells were transduced with lentiCRISPR encoding H1 control or E1A-specific gRNAs, selected for three days, and then stimulated with CT DNA or RIG-I ligand for 24 hours, followed by measuring type I IFN activity in supernatants. *:p<0.05. (B) HEK 293 cells, treated with the indicated ligands, were lysed at the indicated times post transfection, followed by western blot analysis for phosphorylated TBK1 and IRF3. (C) HEK 293 cells and clonal lines of STING KO and TBK1 KO cells were treated with the indicated ligands for 3 hours, followed by western blot for the indicated phosphorylation sites on IRF3. (D) HEK 293 cells and STING KO cells were transduced with lentiCRISPR as described in (A), then stimulated with the indicated ligands for 8 hours followed by western blot analysis.

### A STING-independent DNA sensing pathway (SIDSP) in human cells

To explore these two DNA sensing pathways in more detail, we turned to additional mouse and human cell lines. We first confirmed that the type I IFN response to transfected CT DNA was STING-dependent in primary mouse fibroblasts at the peak of the response four hours post transfection: both DNA- and cGAMP-activated IFN responses were reduced by 99.9% at this time point in STING-deficient fibroblasts (Fig. 2A). We generated two independent clonal lines of STING-deficient human U937 monocytes, a well-characterized human lymphoma cell line (Fig. S2). We transfected these cells with DNA or cGAMP and measured *IFNB1* transcription by quantitative RT-PCR 16 hours after transfection. As expected, both STING-deficient U937 clones failed to respond to cGAMP (Fig. 2B). However, DNA transfection of STING-deficient U937 cells activated a potent type I IFN response (Fig. 2B). We then performed a time course analysis of cGAMP- and DNA-activated IFN responses, comparing control U937 cells to STING-deficient cells. Control U937 cells responded to both DNA and cGAMP transfection with robust *IFNB1* transcription that peaked at 8 hours. STING-deficient U937 cells failed to respond to cGAMP, but they activated a potent antiviral response to DNA that was delayed by several hours, peaking at 16 hours with *IFNB1* mRNA levels that were indistinguishable from those of control cells at this same time point (Fig. 2C). We next tested whether the antiviral response to DNA was dependent on the transcription factors IRF3 and IRF7, which together are essential for the IFN response to all other known nucleic acid detection pathways (*12*). To do this, we used a previously described clonal line of IRF3/IRF7 double knockout human THP1 monocytes (*13*). We found that the potent *IFNB1* transcription in response to both DNA and RNA was completely IRF3/7-dependent at 16 hours post transfection (Fig. 2D). These findings reveal three important points about the DNA-activated antiviral response. First, and consistent with dozens of prior studies, the antiviral response to DNA in mouse cells is nearly entirely STING-dependent (*9*). Second, unlike mouse cells, human cells possess a robust STING-independent DNA sensing pathway that is delayed relative to the cGAS-STING pathway. Third, IRF3 and IRF7 are essential for both DNA-activated antiviral responses in human cells. Thus, human cells – but not mouse cells – have a robust STING-independent DNA sensing pathway (SIDSP).

**Figure 2:**
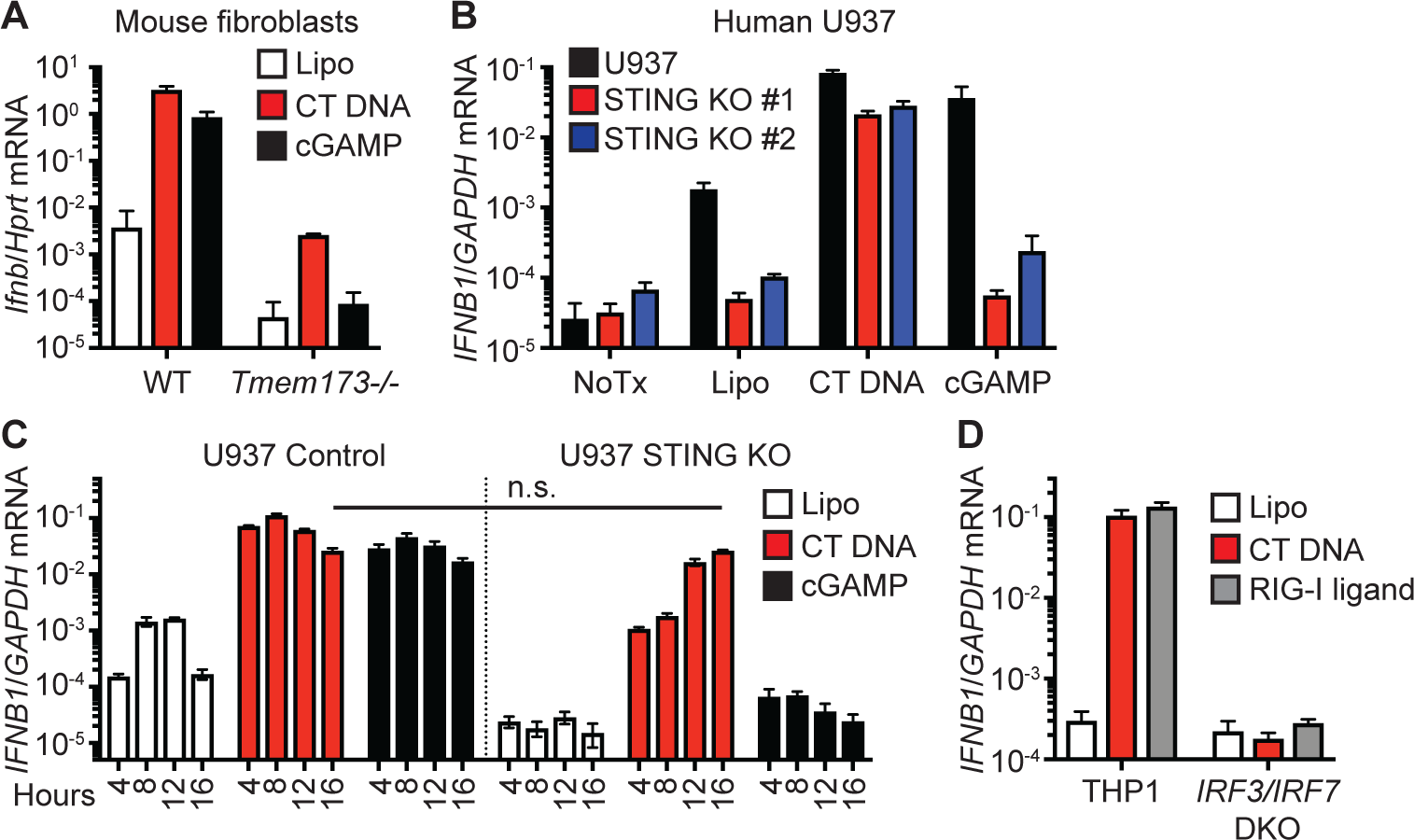
A STING-independent DNA sensing pathway in human cells. (A) Primary mouse embryonic fibroblasts were treated with Lipofectamine alone (Lipo) or the indicated ligands for four hours before harvest and quantitative RT-PCR (qRT-PCR) analysis of *Ifnb* mRNA expression. (B) PMA-differentiated human U937 monocytes or two clonal lines of STING KO U937 cells were treated with the indicated ligands for 16 hours before harvest and qRT-PCR analysis of *IFNB1* mRNA expression. (C) PMA-differentiated WT U937 cells and STING KO U937 cells were treated with the indicated ligands for the indicated times before harvest and qRT-PCR analysis of *IFNB1* mRNA expression, normalized to *GAPDH* expression. n.s.: not significant. (D) PMA-differentiated THP1 cells and IRF3/IRF7 DKO THP1 cells were treated with the indicated ligands for 16 hours before harvest and qRT-PCR analysis of *IFNB1* mRNA expression.

Using our control and STING-deficient U937 cells to genetically separate the cGAS-STING pathway from the SIDSP, we evaluated the structural features of the DNA ligands that triggered these pathways. cGAS activation is mediated by its binding to the sugar phosphate backbone of double-stranded DNA in a sequence-independent manner (*14, 15*). Accordingly, control U937 cells mounted an equally robust antiviral response to both sheared CT DNA and circular plasmid DNA (Fig. 3A). However, STING KO U937 cells responded potently to CT DNA but not to plasmid DNA (Fig. 3A). We found that sonication of the plasmid DNA prior to transfection restored the IFN response in STING KO U937 cells to levels that were identical to those activated by CT DNA (Fig. 3, B and C). Similarly, phosphorylation of IRF3 serine 386 in STING KO HEK 293 cells was potently activated by CT DNA, annealed 100 base-pair DNA oligos (ISD100), and sonicated plasmid DNA, but not by circular plasmid DNA (Fig. 3, D and E). Thus, the cGAS-STING pathway and the SIDSP are activated by different features of DNA: cGAS detects the backbone of dsDNA, whereas the SIDSP detects DNA ends.

**Figure 3:**
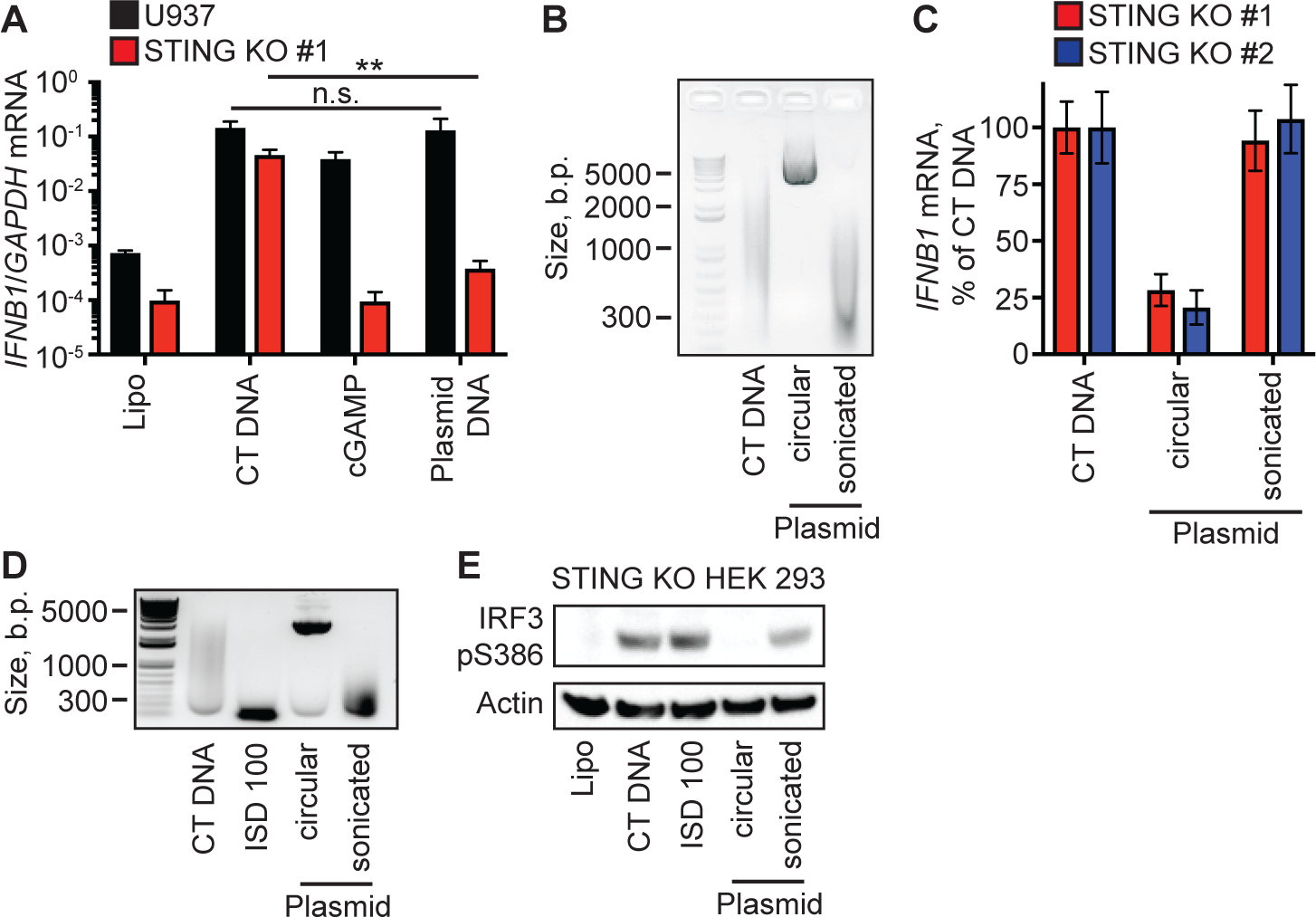
The SIDSP is activated by DNA ends. (A) PMA-differentiated U937 cells and STING KO U937 cells were treated with the indicated ligands for 16 hours before harvest and qRT-PCR analysis of *IFNB1* mRNA expression. n.s.: not significant; **:p<0.01. (B) CT DNA, supercoiled plasmid, and sonicated plasmid were run on a DNA agarose gel and visualized with SYBR-Safe. (C) Two clonal lines of PMA-differentiated STING KO U937 cells were treated with the indicated ligands for 16 hours, with *IFNB1* mRNA expression in CT DNA-treated cells set at 100%. (D) CT DNA, 100 base pair annealed DNA oligos (ISD100), supercoiled and sonicated plasmid DNAs were visualized on DNA-agarose gel. (E) STING KO HEK 293 cells were treated with the indicated ligands for three hours before harvesting lysates and evaluating IRF3 S386 phosphorylation by western blot.

### Human DNA-PK is essential for the SIDSP

Our finding that the activation of the SIDSP requires exposed DNA ends led us to consider two key DNA damage response pathways that are activated by DNA ends: the Ataxia-Telangiectasia Mutated kinase (ATM) pathway that is important for homology-dependent DNA repair, and the DNA-dependent Protein Kinase (DNA-PK) pathway that mediates non-homologous DNA end joining (NHEJ; (*16*). We transfected STING KO U937 cells with CT DNA in the presence of well-characterized chemical inhibitors of the kinase activities of ATM (Ku-60019; (*17*) or DNA-PK (Nu-7441; (*18*). Both of these inhibitors reduced the DNA-activated phosphorylation of the histone H2AX on serine 139 (γ-H2AX) in a concentration-dependent manner, confirming their activity in these cells (Fig. 4A). However, we found that the SIDSP was potently blocked by the DNA-PK inhibitor but unaffected by the ATM inhibitor (Fig. 4B). Moreover, the DNA-PK inhibitor Nu-7441 blocked the DNA-activated antiviral response but had no effect on the RNA-activated RIG-I-MAVS pathway (Fig. 4C).

**Figure 4:**
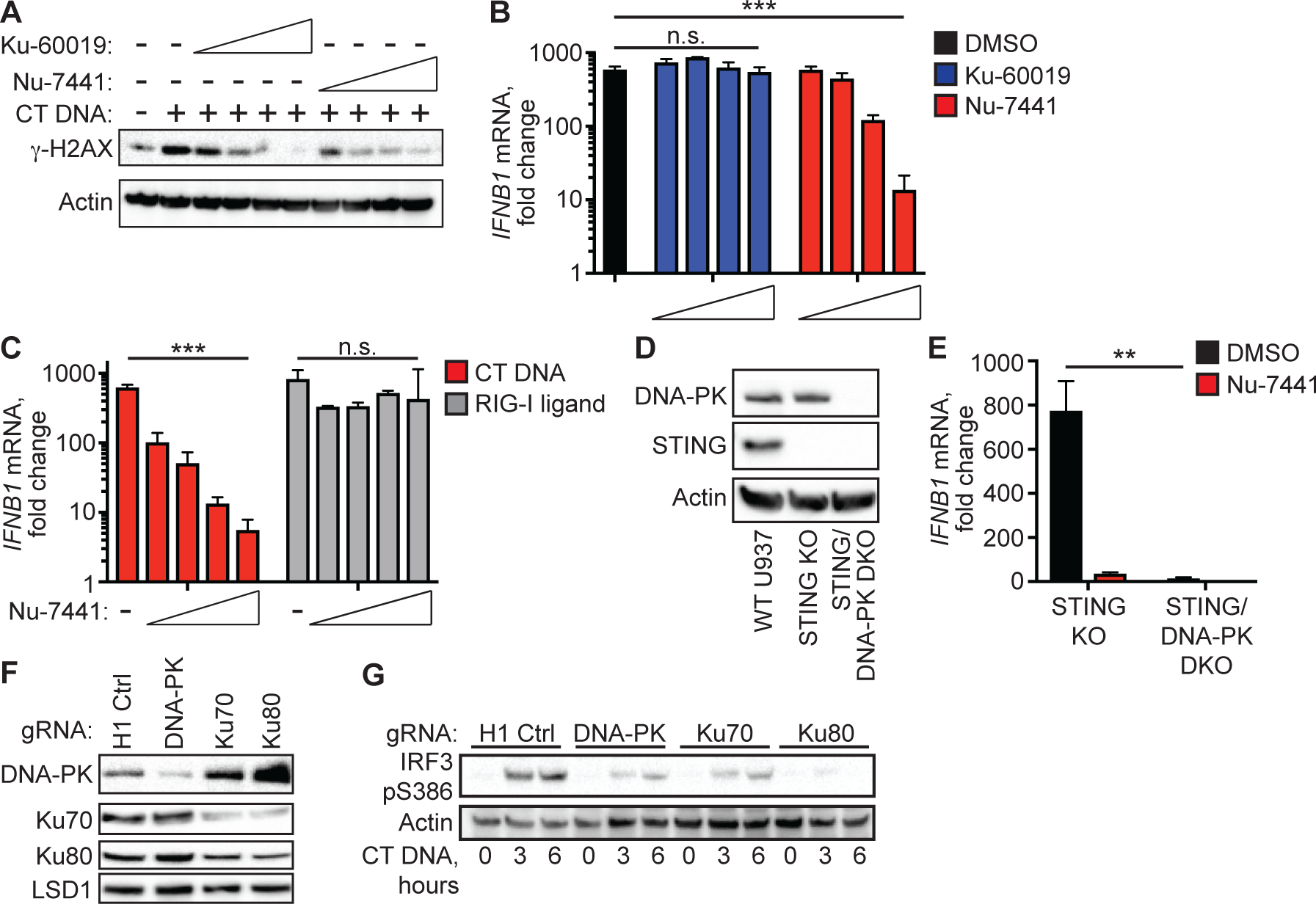
Human DNA-PK is essential for the SIDSP. (A) PMA-differentiated STING KO U937 cells were treated with CT DNA for 16 hours in the presence of DMSO control or increasing concentrations of Ku-60019 ATM inhibitor or Nu-7441 DNA-PK inhibitor, followed by western blot analysis of γ-H2AX phosphorylation. (B) PMA-differentiated STING KO U937 cells were treated with CT DNA for 16 hours in the presence of inhibitors as described in (A), followed by qRT-PCR analysis of *IFNB1* mRNA expression. n.s.: not significant; ***:p<0.001. (C) PMA-differentiated STING KO U937 cells were treated with CT DNA or RIG-I ligand in the presence of DMSO or Nu-7441 for 16 hours, followed by qRT-PCR analysis of *IFNB1* mRNA expression. n.s.: not significant; ***:p<0.001. (D) Western blot analysis of DNA-PK and STING in WT U937 cells and clonal lines of STING KO and STING/D-NA-PK DKO U937 cells. (E) PMA-differentiated STING KO and STING/DNA-PK DKO U937 cells were treated with CT DNA for 16 hours in DMSO or 2μM Nu-7441, followed by qRT-PCR analysis of *IFNB1* mRNA expression, normalized to Lipo control-treated cells. **:p<0.01. (F) STING KO HEK 293 cells were transduced with lentiCRISPR encoding gRNAs specific for the indicated targets, selected for three days in puromycin, and then harvested for western blot analysis of the indicated proteins. (G) STING KO HEK 293 cells from (F) were stimulated with CT DNA for the indicated time points and then harvested for western blot analysis of IRF3 S386 phosphorylation.

We next used lentiCRISPR to simultaneously target U937 cells with guide RNAs targeting STING and the catalytic subunit of DNA-PK (DNA-PKcs), which is encoded by the *PRKDC* gene. DNA-PK-targeted cells were severely compromised for growth relative to control cells, as has been previously reported (*19*), but we managed to generate a clonal line of U937 cells doubly deficient for STING and DNA-PK, verified by western blot and DNA sequencing, together with a third clonal line of STING KO U937 cells (Figs. 4D and S3). Consistent with the chemical inhibitor data, we found that STING/DNA-PK DKO U937 cells were profoundly impaired in their IFN response to DNA (Fig. 4E).

The activation of DNA-PK requires the Ku70 and Ku80 cofactors that are responsible for DNA end binding and recruitment of DNA-PKcs to damaged DNA (*16*). We attempted to generate clonal lines of U937 cells deficient for Ku70 and Ku80 but we were not able to recover live knockout cells, likely because they are essential genes in human somatic cells (*20, 21*). We therefore employed a transient lentiCRISPR approach in STING KO HEK 293 cells to target the *XRCC6* (Ku70) and *XRCC5* (Ku80) genes at the population level. Three days after selection of transduced cells in puromycin, we observed reduced levels of DNA-PKcs, Ku70, and Ku80 proteins in HEK 293 cells targeted with the respective guide RNAs (Fig. 4F). Moreover, targeting Ku70 resulted in reduced expression of Ku80 protein and vice versa, consistent with prior data demonstrating that Ku70 is required for Ku80 protein stability (*22*). We found that IRF3 S386 phosphorylation in response to DNA transfection was reduced in cells targeted for DNA-PKcs, Ku70, and Ku80, suggesting that all three of these components of the DNA-PK complex are essential for SIDSP activation (Fig. 4G). Together, our data provide pharmacological and genetic evidence that DNA-PK is essential for the SIDSP in human cells.

### The DNA-PK SIDSP activates a broad, potent antiviral response

To define the nature of the transcriptional changes in the DNA-PK SIDSP beyond the canonical antiviral cytokine IFNβ, we performed a global mRNA-Seq analysis in WT and STING KO cells, evaluating the changes in gene expression following DNA transfection and the effect of the DNA-PK inhibitor Nu-7441 on this response. After mapping to the human transcriptome, normalizing read counts across all samples, and removing features with fewer than 10 mean counts per million (CPM), our dataset revealed tight concordance among the three biological replicates within each condition and differential clustering of each condition relative to all others (Fig. S4, A and B). A direct comparison of WT and STING KO U937 cells treated with transfection reagent alone revealed largely similar basal gene expression levels, demonstrating that the clonal line of STING KO U937 cells was comparable to control cells (Fig. S4C). Moreover, we found that treatment with Nu-7441 in the absence of DNA stimulation had no significant effect on gene expression in either WT or STING KO cells (Fig. S4, D and E), establishing the baseline conditions used for comparison to the DNA-treated samples within each genotype.

We compared DNA-activated WT and STING KO samples at 8 and 16 hours post-transfection to their respective transfection reagent alone controls, in the presence of DMSO or 2μM Nu-7441. We focused first on the interferon-mediated antiviral response, objectively defined here by compiling genes in this category delineated by Gene Ontology Consortium terms (Table S1). We compiled a list of antiviral response genes with a fold change of greater than 1.5 and a false discovery rate (FDR) of <0.05 in any one of the comparisons. A heat map of these 124 differentially expressed genes revealed a broad, potent, and overlapping antiviral program triggered by DNA in both WT and STING KO cells (Fig. 5A and Table S2). Focusing on the seven IFN genes that were significantly induced in either WT or KO cells, we found that DNA transfection activated broad IFN expression in both WT and STING KO cells (Fig. 5B). Moreover, and consistent with the delayed *IFNB1* response in STING KO cells (Fig. 2C), we found that expression of all IFNs increased between 8 and 16 hours in the STING KO cells, whereas these same genes plateaued or decreased in expression between 8 and 16 hours in WT cells (Fig. 5B). This delay in SIDSP-mediated gene expression held true when objectively comparing all upregulated genes in STING KO and WT cells. Specifically, 912/926 (98.5%) of upregulated genes in STING KO cells continued to increase in gene expression between 8 and 16 hours, with a larger relative increase compared to WT cells (Fig. 5C and Tables S3 and S4). Thus, the SIDSP activates a potent, broad gene expression program that is delayed relative to the DNA-activated antiviral response in WT human cells.

**Figure 5:**
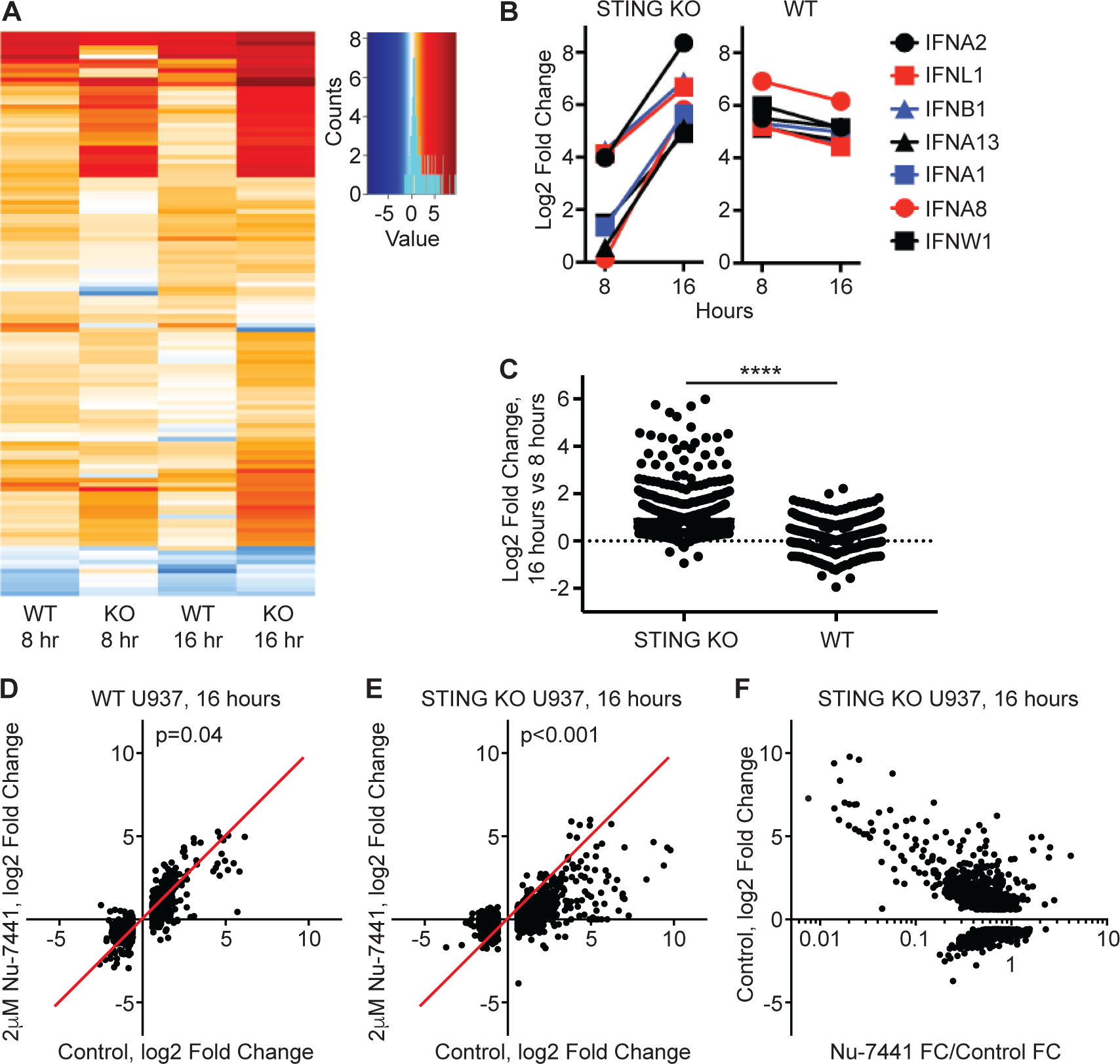
The DNA-PK SIDSP activates a broad gene expression program. (A) Heat map representation of log2 Fold Change in gene expression for the 124 antiviral response genes with signicant differential expression in one of the four comparisons. The key includes a histogram in cyan plotting the distribution of log2 Fold Change values for all the included genes. (B) Expression data for all interferon genes significantly induced in WT or STING KO U937 cells, plotting the log2 Fold Change at 8 and 16 hours post treatment. (C) To measure the trajectories of global gene expression from 8 to 16 hours post CT DNA transfection, the fold change in gene expression at 16 hours was divided by the fold change for the same genes at 8 hours and plotted for all upregulated genes in STING KO (n=926) and WT (n=563). ****:p<0.0001, Mann-Whitney unpaired t-test. (D) In WT U937 cells, the 16 hour CT DNA-activated log2 Fold Change was plotted for DMSO control-treated cells on the x-axis and for Nu-7441-treated cells on the y-axis. Each dot represents a single gene that was differentially expressed at 16 hours in DMSO-treated cells (n=1024). The red line through the origin indicates the line of equivalence representing no effect of the drug treatment. A Wilcoxon matched pairs signed rank test was used to determine the p value between DMSO control and Nu-7441-treated samples after DNA stimulation. (E) In STING KO U937 cells, the effect of Nu-7441 on global gene expression for 1327 genes differentially expressed in DMSO control-treated cells, plotted as in (D). A Wilcoxon matched pairs signed rank test was used to determine the p value between DMSO control and Nu-7441-treated samples after DNA stimulation. (F) In STING KO U937 cells, the log2 Fold Change in DMSO control-treated cells was plotted on the y-axis for differentially expressed genes, and the effect of Nu-7441 on these same genes was plotted in log10 format on the y-axis, n=1327.

We next quantitated the effect of the Nu-7441 DNA-PK inhibitor on global gene expression in WT and STING KO cells. We plotted the fold change values of all differentially expressed genes at 16 hours post DNA transfection, comparing vehicle-treated cells to those treated with 2 μM Nu-7441. In WT cells, we found that Nu-7441 had a mild inhibitory effect on the expression of 718/1024 (70.1%) of differentially expressed genes (Fig. 5D and Table S4), suggesting that at least some of the DNA-activated gene expression in WT human cells reflects the combined contributions of cGAS-STING and the SIDSP. Strikingly, we found that Nu-7441 had an inhibitory effect on 1254/1327 (94.5%) of differentially expressed genes in STING KO cells, including both upregulated and downregulated transcripts (Fig. 5E and Table S3). Moreover, a plot of the fold change in vehicle-treated STING KO cells versus the inhibitory effect of Nu-7441 revealed that the most differentially expressed genes tended to be those most affected by the drug (Fig. 5F and Table S3).

These mRNA-Seq data reveal a number of important features of the DNA-PK SIDSP. First, the SIDSP is a broad and potent antiviral response that results in significant changes in expression of over a thousand human genes. Second, global gene expression in the SIDSP is delayed relative to the DNA-activated antiviral response in WT human cells, highlighting kinetic differences of antiviral signaling that will be interesting to explore in the future. Third, the Nu-7441 inhibitor of DNA-PK kinase activity influences the vast majority of differential gene expression in the SIDSP, as well as a fraction of gene expression in WT cells. Thus, DNA-PK kinase activity is at the apex of the SIDSP, strongly suggesting that it is the primary sensor of this pathway rather than an incidentally activated peripheral component of a distinct pathway. Importantly, these data provide a clear rationale and framework for exploring the utility of DNA-PK inhibitors in IFN-mediated human autoimmune and autoinflammatory disorders.

### Human HSPA8/HSC70 is a target of the DNA-PK SIDSP

In our studies of E1A antagonism of IRF3 phosphorylation, we found that the antibody raised against IRF3 pS386 detected a second protein that was approximately 20 kilodaltons larger than IRF3 in DNA-activated HEK 293 cells (Fig. 6A). We found that this signal was sensitive to phosphatase treatment (Fig. S5A), thus identifying a novel, cross-reactive phosphoprotein that we named “Mystery Protein” (MP). We detected the DNA-activated phosphorylation of MP in all human cell lines examined, including HeLa cells (Fig. 6B), TERT-immortalized human fibroblasts (Fig. 6C), and primary human fibroblasts (Fig. 6D). Three key features of MP matched those that we defined for the SIDSP and led us to study it in more detail. First, MP appeared only in response to DNA, not RIG-I ligand or cGAMP (Fig. 6, A-E). Second, DNA-activated MP phosphorylation was independent of both STING and TBK1 (Fig. 6E). Finally, MP was phosphorylated in response to DNA ends but not circular plasmid DNA (Fig. 6F).

**Figure 6:**
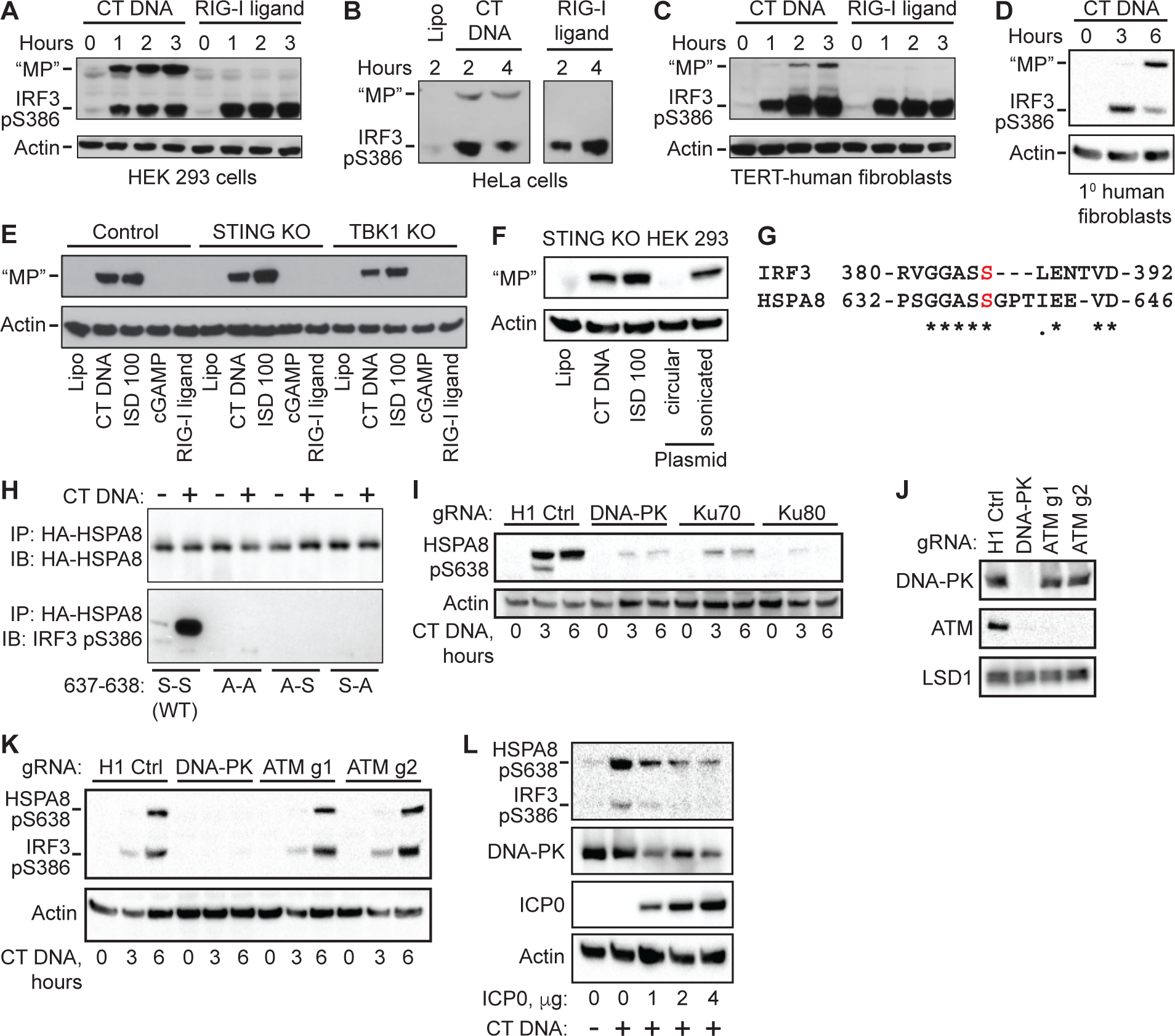
HSPA8 is a target of the DNA-PK-SIDSP. (A-D): The indicated human cells were treated with CT DNA or RIG-I ligand for the indicated tims before harvest and western blot analysis of IRF3 S386 phosphorylation. “Mystery Protein” (”MP”) is indicated on the blots. (E) Control, STING KO, and TBK1 KO HEK 293 cells were treated with the indicated ligands for 3 hours before harvest and western blot analysis of “MP.” (F) STING KO HEK 293 cells were treated with the DNA ligands described in Fig. 3D for 3 hours, followed by western blot analysis of “MP.” (G) Alignments of human IRF3 and HSPA8/HSC70. The red S indicates IRF3 S386 and HSPA8 S638. (H) HEK 293 cells were transfected with plasmids encoding the indicated human HA-HSPA8 constructs, then treated the next day with CT DNA for 3 hours before harvest, HA-immunoprecipitation, and western blot using the IRF3 pS386 antibody. (I) HEK 293 cells targeted for the indicated genes as described in Fig. 4F were treated with DNA and harvested for western blot analysis using the IRF3 pS386 antibody that detects HSPA8 pS638. (J) STING KO HEK 293 cells were transduced with lentiCRISPR targeting H1 control, DNA-PK, or ATM, selected for three days, and then harvested for western blot of the indicated proteins. (K) STING KO HEK 293 cells, transduced and selected as described in (J), were treated with CT DNA and then harvested for western blot analysis of IRF3 S386 and HSPA8 S638 phosphorylation. (L) STING KO HEK 293 cells were transfected with plasmid encoding the ICP0 protein of herpes simplex virus 1. 24 hours later, the cells were stimulated with CT DNA for 3 hours before harvest and western blot analysis of the indicated proteins.

To identify MP, we used the IRF3 pS386 antibody for immunoprecipitation of HEK 293 cell extracts, followed by trypsin digest and mass spectrometry analysis of recovered peptides. To facilitate the identification of MP, we also generated IRF3-deficient HEK 293 cells using lentiCRISPR. Importantly, MP was still robustly phosphorylated after transfection of these IRF3-targeted cells with DNA, demonstrating that MP was not an unusual, slower migrating isoform of IRF3 itself, and that IRF3 was not required for MP phosphorylation (Fig. S5B). We found that the IRF3 pS386 antibody recovered detectable MP from lysates of DNA-transfected cells (Fig. S5B).

Among the peptides identified by mass spectrometry that were specifically enriched by IP with IRF3 pS386 antibody compared to control antibody, one protein in particular caught our attention. Heat shock protein A8 (HSPA8), also known as heat shock cognate 70 (HSC70), matched the predicted mass of MP at ∼73 kilodaltons. Most intriguingly, we noted a sequence at the extreme C terminus of HSPA8 that corresponds precisely to the sequence adjoining S386 in IRF3, suggesting a probable explanation for cross-reactivity of the antibody (Fig. 6G). HSPA8 is an abundant, constitutively expressed member of the heat shock protein (HSP) family of chaperones that participate in the folding of cellular proteins into their native states, either after synthesis on the ribosome or after stress-induced unfolding (*23*).

To test whether MP was HSPA8, we generated expression vectors for hemagglutinin (HA) epitope-tagged human HSPA8 and three mutants in which one or both serines at positions 637 and 638 were mutated to alanines. We transfected each of these constructs into HEK 293 cells, waited 24 hours, and then transfected the cells with CT DNA for three hours before immunoprecipitation of the HA-tagged proteins and blotting for IRF3 pS386. We found that the IRF3 pS386 antibody robustly detected the WT HSPA8 protein after DNA transfection, but it failed to detect the single or double alanine-substituted mutant HSPA8 proteins (Fig. 6H). Thus, the IRF3 pS386 antibody detects phosphorylated human HSPA8, and both serines 637 and 638 of HSPA8 are essential for antibody binding. Because the IRF3 pS386 antibody was raised against a phosphopeptide in which only S386 was phosphorylated, and because S638 of HSPA8 aligns with S386 of IRF3, we suggest that HSPA8 is phosphorylated, at minimum, on S638 in response to DNA detection. Importantly, IRF3 is also known to be phosphorylated on S385 upon activation (*7*), so it is possible that HSPA8 is additionally phosphorylated on S637.

Similar to the data presented for IRF3 (Fig. 4F), we found that DNA-PK, Ku70, and Ku80 were all essential for the robust phosphorylation of HSPA8 on S638 (Fig. 4I). Next, we compared the effect of DNA-PK and ATM disruption on phosphorylation of IRF3 and HSPA8. We confirmed previous findings that targeting DNA-PK resulted in a loss of ATM protein expression (*24*), and we found that disruption of ATM did not affect DNA-PK protein levels (Fig. 6J). Consistent with the pharmacological data in U937 monocytes (Fig. 4B), we observed that DNA-PK was essential for both IRF3 and HSPA8 phosphorylation after DNA transfection in HEK 293 cells, but loss of ATM had no effect on this response (Fig. 6K). Finally, we evaluated the effect of the ICP0 ubiquitin ligase of herpes simplex virus 1 (HSV-1), which targets DNA-PK for degradation (*25*), on activation of the SIDSP. We found that ICP0 expression in STING KO HEK 293 cells blocked DNA-activated phosphorylation of IRF3 and HSPA8 in a dose-dependent manner, revealing a second viral antagonist of the SIDSP (Fig. 6L). Thus, we have identified phosphorylation of human HSPA8 on serine 638 as a novel target of the DNA-PK SIDSP. Phosphorylation of the C-terminus of HSPA8 has been proposed to modulate its interactions with cochaperone proteins (*26*), but inducible, site-specific phosphorylation of HSPA8 has not been reported previously.

### HSPA8 delineates the antiviral modality of human DNA-PK

We noted that the amino acids surrounding serine 638 of HSPA8 are completely conserved across mammalian evolution, unlike those surrounding IRF3 (Fig. S5C). We took advantage of this conservation to test whether we could detect activation of the DNA-PK SIDSP in cell lines from primates and rodents, using HSPA8 phosphorylation as a convenient marker. We found that HSPA8 phosphorylation was robustly activated after DNA transfection in cells from all primate species tested and in rat cells, but that mouse fibroblasts failed to activate HSPA8 phosphorylation (Figs. 7A and S5D). We then introduced epitope-tagged mouse HSPA8 into human HEK 293 cells, and human HSPA8 into mouse fibroblasts. Mouse HSPA8 was robustly phosphorylated in human cells, but neither human nor mouse HSPA8 became phosphorylated in mouse fibroblasts after DNA transfection (Fig. 7, B and C). Together with the well documented observation that nearly all of the IFN response to DNA in mouse cells is STING-dependent (Fig. 2A), our data suggest that the DNA-PK SIDSP is present in humans, primates, and rats, but absent or severely impaired in mouse cells.

**Figure 7:**
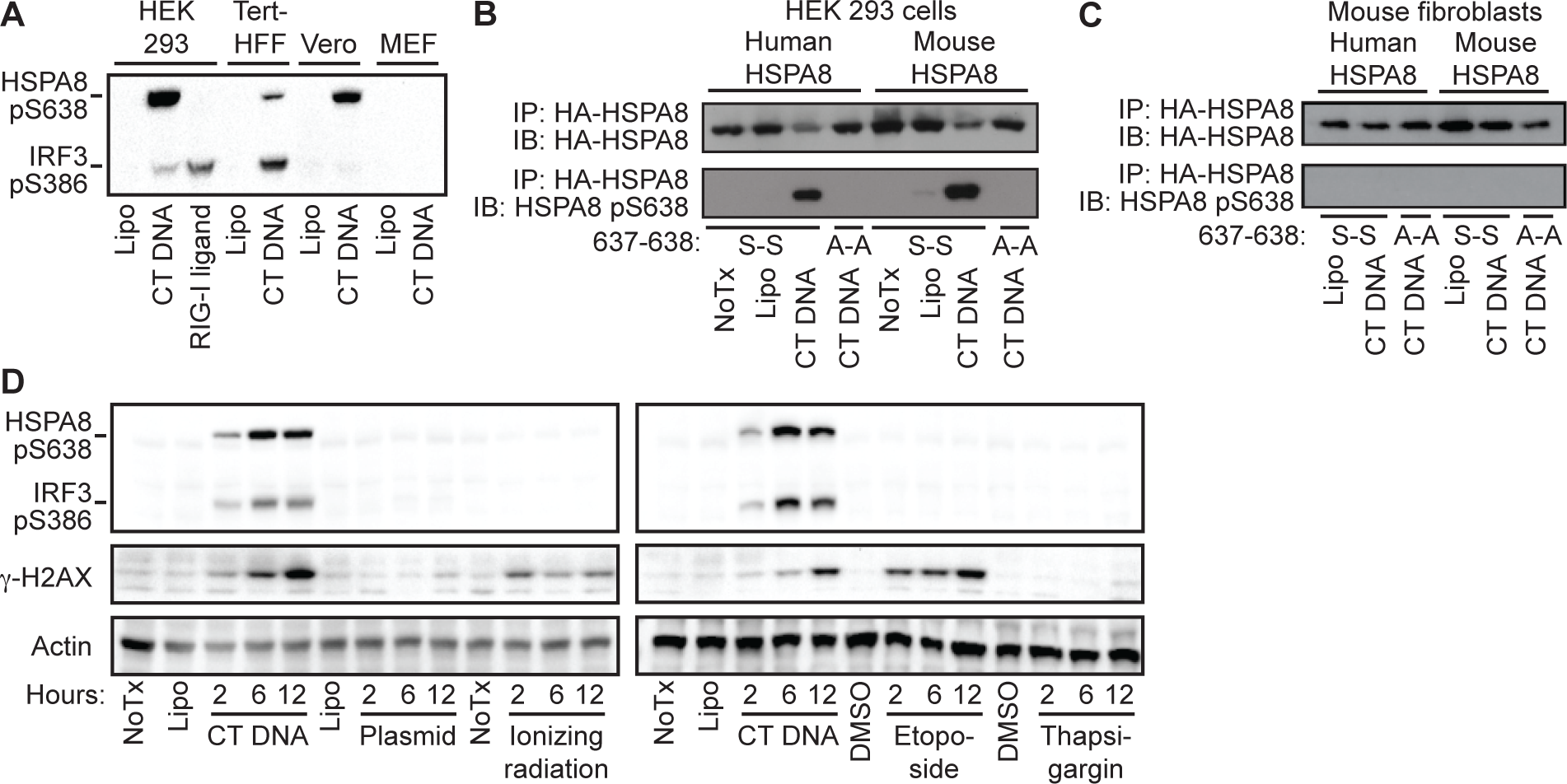
HSPA8 delineates the antiviral modality of human DNA-PK. (A) The indicated human, primate, and mouse cell lines were stimulated with CT-DNA for 3 hours before harvest and western blot for the indicated proteins. (B) HEK 293 cells were transfected with either human HA-HSPA8 constructs or mouse HA-HSPA8 constructs, followed by CT DNA stimulation for 3 hours, HA immunoprecipitation, and western blot analysis of the indicated proteins. (C) Mouse immortalized Jackson fibroblasts were transfected and treated as indicated in (B). (D) HEK 293 cells were stimulated with CT DNA or supercoiled plasmid DNA, or treated with 30 Gray ionizing γ-irradiation, 50μM Etoposide, or 500 nM Thapsigargin before harvest at the indicated time points and western blot analysis of IRF3 pS386, HSPA8 pS638, and γ-H2AX.

We next tested for activation of the SIDSP in response to DNA damage, which potently triggers activation of DNA-PK (*16*). We treated STING KO HEK 293 cells with CT DNA, plasmid DNA, ionizing radiation, or the topoisomerase-II inhibitor etoposide, monitoring activation of both IRF3 and HSPA8 phosphorylation up to 12 hours after treatment. As shown in Figs. 3E, 4A, and 6F, we found that CT DNA, but not circular plasmid DNA, induced robust phosphorylation of both IRF3 and HSPA8, together with potent activation of H2AX S139 phosphorylation (Fig. 7D). However, and intriguingly, neither ionizing radiation nor etoposide activated IRF3 or HSPA8 phosphorylation, despite robust H2AX S139 phosphorylation and the well characterized activation of DNA-PK by each of these DNA damaging agents (Fig. 7D; (*16*). As a control, we used thapsigargin, which induced a potent endoplasmic reticulum (ER) stress response, but not IRF3, HSPA8, or H2AX phosphorylation (Fig. 7D). Together, our data suggest two distinct modalities of DNA-PK activation. First, the well-characterized role of DNA-PK in response to DNA damage, which involves coordination of the NHEJ repair machinery, does not activate DNA-PK-dependent phosphorylation of IRF3 or HSPA8. Second, the DNA-PK-dependent response to foreign DNA triggers both IRF3 and HSPA8 phosphorylation. Thus, the SIDSP activates unique targets of DNA-PK in a manner that is distinct from that triggered by DNA damage.

## Discussion

We have identified DNA-PK as the sensor of a potent, STING-independent DNA sensing pathway (SIDSP) that is present in human cells but weak or absent from mouse cells. We identify two DNA virus-encoded antagonists of the DNA-PK SIDSP, and we show that a small molecule inhibitor of DNA-PK kinase activity potently reduces the robust and broad transcriptional response triggered by foreign DNA in human cells. Finally, we present evidence that the DNA-PK SIDSP includes unique targets that are triggered only by foreign DNA and not by DNA damage. The existence of a second DNA sensing pathway that is present in human cells but not mouse cells has important implications for our understanding of antiviral immunity, for ongoing efforts to inhibit the antiviral response to treat autoimmune diseases, and for the possibility of harnessing this pathway to enhance immune responses to tumors.

Prior studies have suggested a role for DNA-PK in the antiviral response triggered by detection of intracellular DNA (*27–29*). However, our findings differ from these studies in three important ways. First, DNA-PK was proposed to act upstream of STING (*27*), whereas we use clonal lines of STING-deficient human cells to demonstrate that DNA-PK triggers a STING-independent antiviral pathway. Second, the contribution of DNA-PK in prior studies was found to be independent of its catalytic activity (*27*). We show that the vast majority of differential gene expression in the SIDSP in human cells is sensitive to a chemical inhibitor of DNA-PK kinase activity. Finally, DNA-PK was suggested to contribute to the DNA-activated IFN response in mouse cells (*27–29*). We previously found that Ku70- and DNA-PKcs-deficient mouse cells had no defect in DNA-activated IFN production (*30*), and we find no evidence here for activation of the DNA-PK SIDSP in mouse cells. Despite these important differences in the signaling requirements and species specificity of DNA-PK activity, we note that these studies identified multiple proteins from Vaccinia virus that inhibit DNA-PK activation (*28, 29*). Moreover, interactions between hAd5 E1A and Ku70, as well as between hAd5 E4 proteins and DNA-PK, have been described previously (*31, 32*). Together with our identification of hAd5 E1A and HSV-1 ICP0 as potent inhibitors of the DNA-PK SIDSP, it is clear that numerous classes of DNA viruses encode DNA-PK antagonists.

We found that the Nu-7441 DNA-PK inhibitor potently reduced nearly all gene expression triggered by the SIDSP, demonstrating that DNA-PK kinase activity drives the SIDSP transcriptional response. A key question for future studies is whether DNA-PK directly phosphorylates IRF3 and HSPA8, or whether another kinase downstream of DNA-PK is required. DNA-PK was previously shown to directly phosphorylate human IRF3 on threonine 135 (*33*); this site matches the known preferred DNA-PK phosphorylation motif of a serine or threonine followed by a glutamine (*16*). However, neither of the phosphorylation sites that we identified as DNA-PK-dependent in HSPA8 or IRF3 match this motif, suggesting that if DNA-PK directly phosphorylates these sites, it does so in a non-canonical manner.

We identified serine 638 of HSPA8 as a unique and specific target of the DNA-PK SIDSP in human cells. Intriguingly, the original description of DNA-PK kinase activity over 30 years ago found inducible phosphorylation of a distinct heat shock protein, HSP90, in extracts of human cells but not mouse cells (*34*). We are currently exploring the role of HSPA8 and its phosphorylation in the SIDSP, which has been complicated thus far by the essential role that HSPA8 plays in cellular homeostasis and survival (*23*). However, we used the conservation of HSPA8 among mammals as a means to explore the activation of the DNA-PK SIDSP in primates and rodents. Consistent with the lack of a significant STING-independent IFN response in mouse fibroblasts, we found that HSPA8 phosphorylation did not occur in mouse cells. However, all primates tested, as well as rats, demonstrated intact HSPA8 phosphorylation, suggesting that the SIDSP is broadly present in mammals and that laboratory mice specifically lost a robust SIDSP after their divergence from the common ancestor of mice and rats. The underlying reason for the weak or absent SIDSP in mice is a very interesting question for future studies, the goal of which will be to restore the SIDSP in mice and then explore its role in antiviral immunity *in vivo*.

We found that DNA damage and the SIDSP trigger distinct outcomes of DNA-PK kinase activity, with only foreign DNA triggering HSPA8 and IRF3 phosphorylation and the antiviral response. Thus, the same kinase, activated by DNA ends, targets distinct substrates depending on the context of the stimulus. What underlies these two modalities of DNA-PK activity and how are they enforced? Potential explanations include different subcellular localization of foreign DNA versus damaged genomic DNA, as well as accessory proteins that may target DNA-PK to its distinct substrates. It will be important to further define and understand these modalities of DNA-PK in molecular detail. Such an understanding would enable studies of whether these modalities can be overridden, for example converting DNA damage into a potent antiviral response in tumors.

The cGAS-STING antiviral response has become the subject of intense development in the pharmaceutical industry, including efforts to develop inhibitors of cGAS and STING to treat human autoimmune diseases (*35–37*), as well as agonists of STING to improve immune responses to tumors (*38–40*). Our discovery of a second DNA-activated antiviral response in human cells has important implications for these efforts. First, if both cGAS-STING and the DNA-PK SIDSP are activated by intracellular DNA in human autoimmune disease, it is possible that cGAS inhibitors will need to be paired with DNA-PK inhibitors to have therapeutic effect. Importantly, highly specific and efficacious inhibitors of DNA-PK already exist, whereas effective cGAS inhibitors are still being developed. Second, the possibility of harnessing agonism of the DNA-PK SIDSP to trigger innate immune responses in the tumor microenvironment could broaden the toolkit of sophisticated adjuvant immunotherapies.

In summary, we have described the existence of a potent STING-independent DNA sensing pathway (SIDSP) in human cells, and we have identified its sensor, a unique target, two distinct viral antagonists, and a potent small molecule inhibitor of the response. Further study of this new pathway, its relevance in human disease, and its relationship to the cGAS-STING pathway will uncover new insights into human innate immunity.

## Methods

### Reagents, antibodies, and inhibitors

A separate table of the source of all reagents, antibodies, kits, and inhibitors is included with the manuscript. Sheared CT DNA (Sigma) and 2’3’ cGAMP (Invivogen) were purchased and diluted in water; ISD oligos were ordered from Integrated DNA Technologies and annealed in water (*30*); RIG-I ligand was synthesized *in vitro* as previously described using HiScribe T7 High Yield RNA Synthesis Kit (*41*). For plasmid stimulations, midiprepped pcDNA3 was either untreated or sonicated with a Covaris M220 focused ultrasonicator at 5% ChIP (factory setting). Nu-7441 and Ku-60019 (SelleckChem) were suspended in DMSO and used to treat cells for 1 hour prior to stimulation with nucleic acid ligands. For Nu-7441, we used 0.25, 0.5, 1, or 2 μM. For Ku-60019, we used 0.125, 0.25, 0.5, or 1 μM. Untreated cells received the same amounts of plain DMSO.

### Cell treatments

HEK 293 cells were grown in DMEM supplemented with 10% FCS, L-glutamine, penicillin/streptomycin, sodium pyruvate, and HEPES. U937 and THP1 cell lines were grown in RPMI supplemented as above, and differentiated prior to stimulation using 100nM phorbol myristoyl acetate (PMA) for 24 hours and then rested in media lacking PMA for 24 hours.

HEK 293 cells were plated at 0.5 million/well in a 6 well dish in 2mL media the day before stimulation for protein harvest. For RNA harvest and qPCR, U937 cells were plated at 0.25 million/well in a 24 well dish. In the 6 well dish format, cells received 8 μg of CT DNA, ISD100, or pcDNA3 complexed with 8 μl of Lipofectamine 2000. 10 μM cGAMP was complexed with 8 μl Lipofectamine and 1 μg RIG-I ligand was complexed with 1 μl Lipofectamine to achieve comparable induction of IFN across treatments in competent cells. Stimulations done in 24 well plates were scaled by ¼. Etoposide (prepared in DMSO) was diluted in culture media to 50 μM, and untreated cells received the same volume of DMSO. Cells were irradiated with 30 Gy using a Rad Source RS 2000 X-irradiator.

### IFN bioassay

Supernatants from stimulated cells were harvested 24 hours post-stimulation and used to stimulate a HeLa cell line stably expressing an ISRE-luciferase reporter as described previously (*11*).

### Western blotting

Cells were harvested by trypsinization (U937 cells) or vigorous wash with PBS (HEK293 cells), pelleted, and lysed using either a 1% Triton-X-100 buffer (20 mM HEPES, 150 mM NaCl, 10% glycerol, 1 mM EDTA, Pierce phosphatase/protease inhibitors) or, for samples requiring measurement of DNA-PK protein levels, RIPA buffer (150 mM NaCl, 1% Triton-X-100, 0.5% sodium deoxycholate, 0.1% SDS, 50 mM Tris pH 8.0, Pierce phosphatase/protease inhibitors). Lysates were vortexed and incubated on ice for 15 minutes before clearing by centrifugation for 15 minutes. Proteins were separated on Bolt 4–12% Bis-Tris gels (ThermoFisher) in MES buffer for 30 minutes at 200 V and transferred to Immobilon-FL PVDF membrane (Sigma). Blots were blocked in 5% BSA/TBST for 30 minutes prior to incubation with primary. The pIRF3 S386 blots were incubated at 4°C overnight and washed at least 30 minutes in TBST prior to secondary incubation to prevent background. In order to better resolve DNA-PK (470 kDa), lysates were run on 3-8% Tris-acetate gels (ThermoFisher) for 2 hours at 150 V and then transferred in 5% methanol for 3 hours at 20 V at 4°C.

### LentiCRISPR targeting

VSV-G pseudotyped, self-inactivating lentivirus was prepared by transfecting a 60-80% confluent 10-cm plate of HEK 293T cells with 1.5 μg of pVSV-G expression vector, 3 μg of pMDLg/pRRE, 3 μg pRSV-Rev and 6 μg of pRRL lentiCRISPR vectors using Poly(ethyleneimine) (PEI; Sigma). Media was replaced 24 hours post-transfection and harvested 24 hours later for filtration with a 0.45 μm filter (SteriFlip, Millipore). Approximately 1 million cells were transduced with 10 mL filtered virus. Targeting NHEJ components efficiently was difficult; best results were achieved by increasing transduction rates with sequential transductions on two consecutive days. Cells were plated for stimulations while still under selection at day 4 post first transduction.

For CRISPR/Cas9 gene targeting, we generated pRRL lentiviral vectors in which a U6 promoter drives expression of a gRNA, and an MND promoter drives expression of Cas9, a T2A peptide, and either a puromycin or blasticidin (*42*). gRNA sequences are as follows, where the (G) denotes a nucleotide added to enable robust transcription off the U6 promoter and the underlined sequence denotes the Protospacer Adjacent Motif (PAM): H1 off-target control: (G)ACGGAGGCTAAGCGTCGCAA (*43*); *TMEM173* (STING): GGTGCCTGATAACCTGAGTA<u>TGG</u> (*42*); *TBK1:* (G)CATAAGCTTCCTTCGTCCAG<u>TGG</u> (*7*); *PRKDC (*DNA-PK): GCAGGAGACCTTGTCCGCTG<u>CGG</u>; *XRCC6* (Ku70): GATCCGTGGCCCATCATGTCT<u>TGG</u>; *XRCC5* (Ku80): GTTGTGCTGTGTATGGACGT<u>GGG</u>; *ATM* guide 1: (G)CCAAGGCTATTCAGTGTGCG<u>AGG</u> (*43*); *ATM* guide 2: (G)TGATAGAGCTACAGAACGAA<u>AGG</u> (*43*); and E1A (G)AAGACCTGCAACCGTGCCCG<u>GGG</u> (Lau, et al 2015) (*11*). Guides against *PRKDC, XRCC6,* and *XRCC5* were designed using Benchling.

### Generation of clonal cell lines

KO cell lines were generated by limiting dilution, screened by western blot, and verified by Sanger sequencing and functional assays. The STING/DNA-PK DKO U937 cell line was produced by transducing U937s simultaneously with a STING lentiCRISPR puro virus and a DNA-PK lentiCRISPR blasticidin virus, selecting in 10 μg/ml puro and 5 μg/ml blasticidin, and seeding in 96 well plates immediately after selection. Very few colonies grew, and the verified DKO clone grew markedly slower than H1 non-targeted control clones or the STING KO clones, as expected (*19*).

PCR primers used for amplifying genomic DNA surrounding CRISPR targeting sites in clonal lines were as follows (Forward/Reverse): *TMEM173*: 5’-AGCTCCAGGCCCGGATTCG-3’/5’-TGCCCGTTCTCCAGAAGCTC-3’ *TBK1*: 5’-CCCTACTGTATCCTCATG-3’/5’-CTTACCTCCTCTTCAATAGC-3’ *PRKDC:* 5’-GGGGCATTTCCGGGTCCGGG-3’/5’-TGCCCTGCCCCCCACTCTGC-3’ Amplicons were cloned using the Zero Blunt TOPO PCR Cloning kit (ThermoFisher), prepared as plasmids, and then several individual plasmids were sequenced. Sequencing alignments were made using Benchling.

### RNA isolation and qPCR

Cells were harvested in Trizol before purification via Direct-zol RNA miniprep (Genesee Scientific) per manufacturer’s instructions with an additional dry spin after disposing of the final wash to prevent carryover. cDNA was generated using EcoDry double primed premix (Clontech). qPCR was performed using iTaq supermix on the Bio-Rad CFX96 Real-Time system. Human gene PCR primer sequences are as follows:

*GAPDH* Fwd: 5’-AACAGCCTCAAGATCATCAGC-3’, *GAPDH* Rev: 5’-CACCACCTTCTTGATGTCATC-3’

*IFNB1* Fwd: 5’-ACGCCGCATTGACCATCTATG-3’, *IFNB1* Rev: 5’-CGGAGGTAACCTGTAAGTCTGT-3’.

Mouse primer sequences are as follows:

*Hprt* Fwd: 5’-GTTGGATACAGGCCAGATTTGTTG-3’, *Hprt* Rev: 5’-GAGGGTAGGCTGGCCTATAGGCT-3’

*Ifnb* Fwd: 5’-GCACTGGGTGGAATGAGACTATTG-3’ *Ifnb Rev: 5’*-TTCTGAGGCATCAACTGACAGGTC-3’.

### mRNA-Seq and analysis

Total RNA was added directly to lysis buffer from the SMART-Seq v4 Ultra Low Input RNA Kit for Sequencing (Takara), and reverse transcription was performed followed by PCR amplification to generate full length amplified cDNA. Sequencing libraries were constructed using the NexteraXT DNA sample preparation kit (Illumina) to generate Illumina-compatible barcoded libraries. Libraries were pooled and quantified using a Qubit® Fluorometer (Life Technologies). Dual-index, single-read sequencing of pooled libraries was carried out on a HiSeq2500 sequencer (Illumina) with 58-base reads, using HiSeq v4 Cluster and SBS kits (Illumina) with a target depth of 5 million reads per sample. Base calls were processed to FASTQs on BaseSpace (Illumina), and a base call quality trimming step was applied to remove low-confidence base calls from the ends of reads. The FASTQs were aligned to the human reference genome using the STAR aligner, and gene counts were generated using htseq-count. QC and metrics analyses were performed using the Picard family of tools (v1.134).

Exploratory analysis and statistics were run using R (version 3.5.1) and bioconductor (version 3.7). The gene count matrix was filtered by a row mean of ten or greater counts and normalized with EDGER. LogCPM transformation was performed using voom through the limma bioconductor package (3.34.8). Statistical analysis (including differential expression) was performed using the limma package (*44, 45*).

Co-expression was performed on genes statistically significant in the differential expression analysis (threshold: linear fold change >= | 1.5 | and FDR <= 0.05) in at least one comparison. The union of these DE genes were loaded into R and filtered by known interferon signaling genes using all of the GO terms listed in Table S1. Correlations (ward.2 clustering and euclidean distance) were run on the union of log2FC using the WGCNA and heatmap.2 bioconductor packages in R (*44, 46, 47*).

### Immunoprecipitation and Mass Spectrometry

To identify HSPA8 using mass spectrometry, we performed immunoprecipitation of CT-DNA stimulated HEK293 cells using the antibody to IRF3 pS386 crosslinked to Dynabeads (ThermoFisher) overnight at 4 C, then washed three times in lysis buffer and two times in ammonium biocarbonate (50mM) before peptide digestion (V5280, Promega). Peptides were loaded onto a 3-cm self-packed C18 capillary pre-column (Reprosil 5 μM, Dr. Maisch). After a 10-min rinse (0.1% Formic Acid), the pre-column was connected to a 25-cm self-packed C18 (Reliasil 3 μM, Orochem) analytical capillary column (inner diameter, 50-μm; outer diameter, 360-μm) with an integrated electrospray tip (∼1-μm orifice). Online peptide separation followed by mass spectrometric analyses was performed on a 2D-nanoLC system (nanoAcquity UPLC system, Waters Corp.). Peptides were eluted using a 150-min gradient with solvent A (H_2_O/Formic Acid, 99.9:1 (v/v)) and B (Acetonitrile/Formic Acid, 99.9:1 (v/v)): 10 min from 0% to 10% B, 105 min from 10% to 40% B, 15 min from 40% to 80% B, and 20 minutes with 100% A. Eluted peptides were directly electrosprayed into a Orbitrap QExactive mass spectrometer (Thermo Fisher Scientific) equipped with a high energy collision cell (HCD). The mass spectrometer was operated in a data-dependent mode to automatically switch between MS and MS/MS acquisitions. Each full scan (from *m/z* 300-1500) was acquired in the Orbitrap analyzer (resolution = 70,000), followed by MS/MS analyses on the top twenty most intense precursor ions that had charge states greater than one. The HCD MS/MS scans were acquired using the Orbitrap system (resolution = 17,500) at normalized collision energy of 28%. The precursor isolation width was set at 2 *m/z* for each MS/MS scan and the maximum ion accumulation times were as follows: MS (100ms), MS/MS (100ms). MS/MS data files were searched using the Comet algorithm (*48*), and the data were further processed using the Institute for System’s Biology’s Trans-Proteomic Pipeline (*49*). Static modification of cysteine (carbamidomethylation; 57.02 Da) was used in the search.

### Cloning of HSPA8

PCR and InFusion cloning (Clonetech) were used to generate N-terminal HA-tagged WT and alanine mutant human HSPA8 constructs from HEK 293 cell cDNA. A murine HSPA8 cDNA clone (Transomic technologies, Clone ID BC089322) was used as template to generate the epitope-tagged mouse versions.

### Transient expression of ICP0

0.25 million STING KO HEK293s were seeded in 12 well format the day before transfection with 0, 1, 2 or 4 μg of ICP0 expression plasmid using Lipofectamine 2000 at a 1 μl:1 μg DNA ratio. Empty pcDNA3 was used to bring the total amount of transfected DNA up to 4 μg total. 24 hours post-transfection, the cells were treated with 4 μg CT DNA or 4 μL Lipofectamine 2000 alone and harvested 3 hours later in RIPA buffer with phosphatase inhibitors for analysis by western blot.

### Experimental replicates and statistics

All experiments presented in this study, except the mRNA-Seq studies, were done two or more times, with biological triplicates for each condition in RT-qPCR experiments. Quantitative data were visualized and analyzed using GraphPad Prism software. Multiple unpaired t-tests with significance determined by Holm-Sidak method were used to compare differences between groups, unless otherwise noted for specific tests in figure legends. Significance is indicated as follows: *p<0.05. **p < 0.01, ***p<0.001, ****p < 0.0001.

## Acknowledgements

We are grateful to Vivian Gersuk and the Benaroya Research Institute Genomics Core for mRNA-Seq, and to all the members of the Stetson lab for discussions.

## Author Contributions

K.B. and D.B.S. conceived and designed the study. K.B., J.H.M., and S.C. performed experiments. S.C. and R.G. analyzed mRNA-Seq data. R.C.J. performed mass spectrometry and analysis. M.G. Jr. provided key resources and data analysis. K.B., J.H.M., S.C., R.G., R.C.J., and D.B.S. wrote and/or edited the manuscript.

---

**Figure S1:**
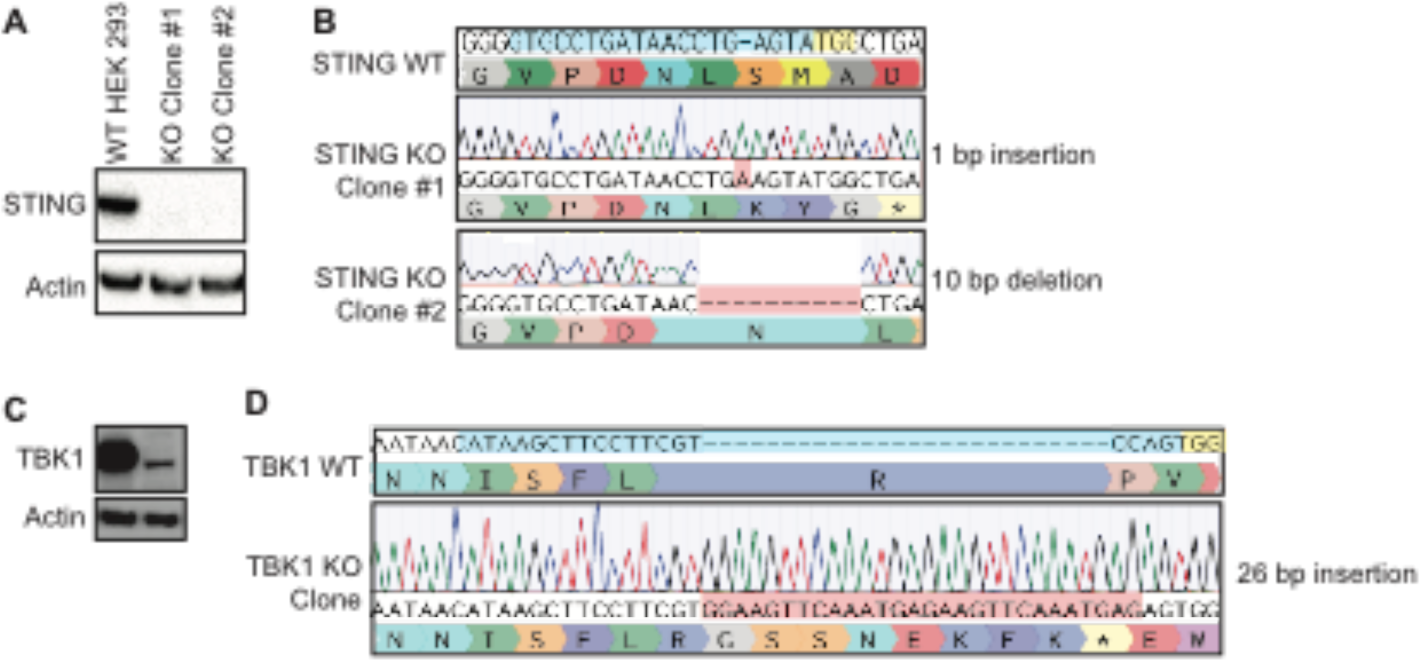
Validation of STING KO and TBK1 KO clonal HEK 293 cell lines. (**A**) HEK 293 cells were transduced with lendCRISPR encodng gRNAs specific for *TMEM173* (STING), selected, cloned by limiting dilution, and then evaluated for STING protein expression by western blot. (**B**) Clonal lines of HEK 293 cells were evaluated by Sanger sequencing revealing a homozygous 1 bp insertion in Clone #1 and a homozygous 10 bp deletion In Clone #2. (**C**) HEK 293 cells wore transduced wflh lentiCRISPR encoding gRNAs specific for TBK1. selected, cloned by muting dilution, and then evaluated for *TBK1* proton expression by western blot. (**D**) The clonal Iine of TBK1-deficient HEK 293 cells was evaluated by Sanger sequencng. revealng a homozygous 26 bp Insertion.

**Figure S2:**
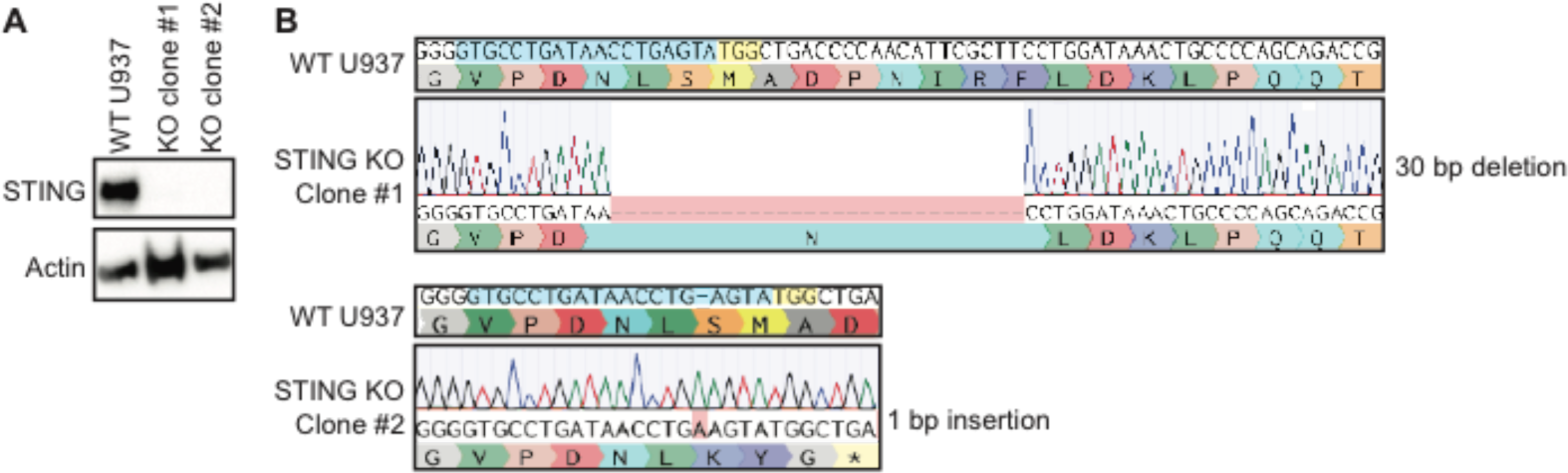
Validation of STING KO clonal U937 cell lines. (**A**) U937 cells were transduced with lentiCRISPR encoding gRNAs specific for *TMEM173* (STING), selected, cloned by limiting dilution, and then evaluated for STING protein expression by western blot. (**B**) Clonal lines of U937 cells were evaluated by Sanger sequencing, revealing a homozygous 30 bp deletion in Clone #1 and a homozygous 1 bp deletion in Clone #2.

**Figure S3:**
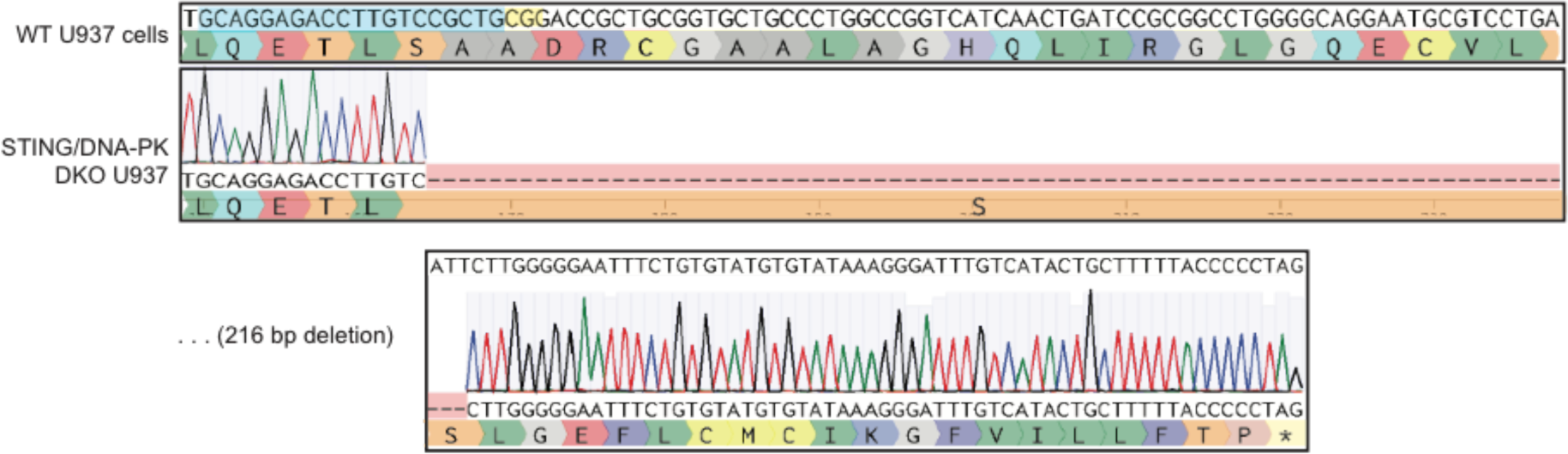
Sequencing of STING/DNA-PK DKO clonal U937 cell line. U937 cells were transduced with lentiCRISPR encoding gRNAs specific for *TMEM173* (STING) and *PRKDC* (DNA-PK), selected, cloned by limiting dilution, and then evaluated by Sanger sequencing of the DNA surrounding the DNA-PK target site, revealing a homozygous 216 bp deletion that entered the following intron, disrupting the splice site.

**Figure S4:**
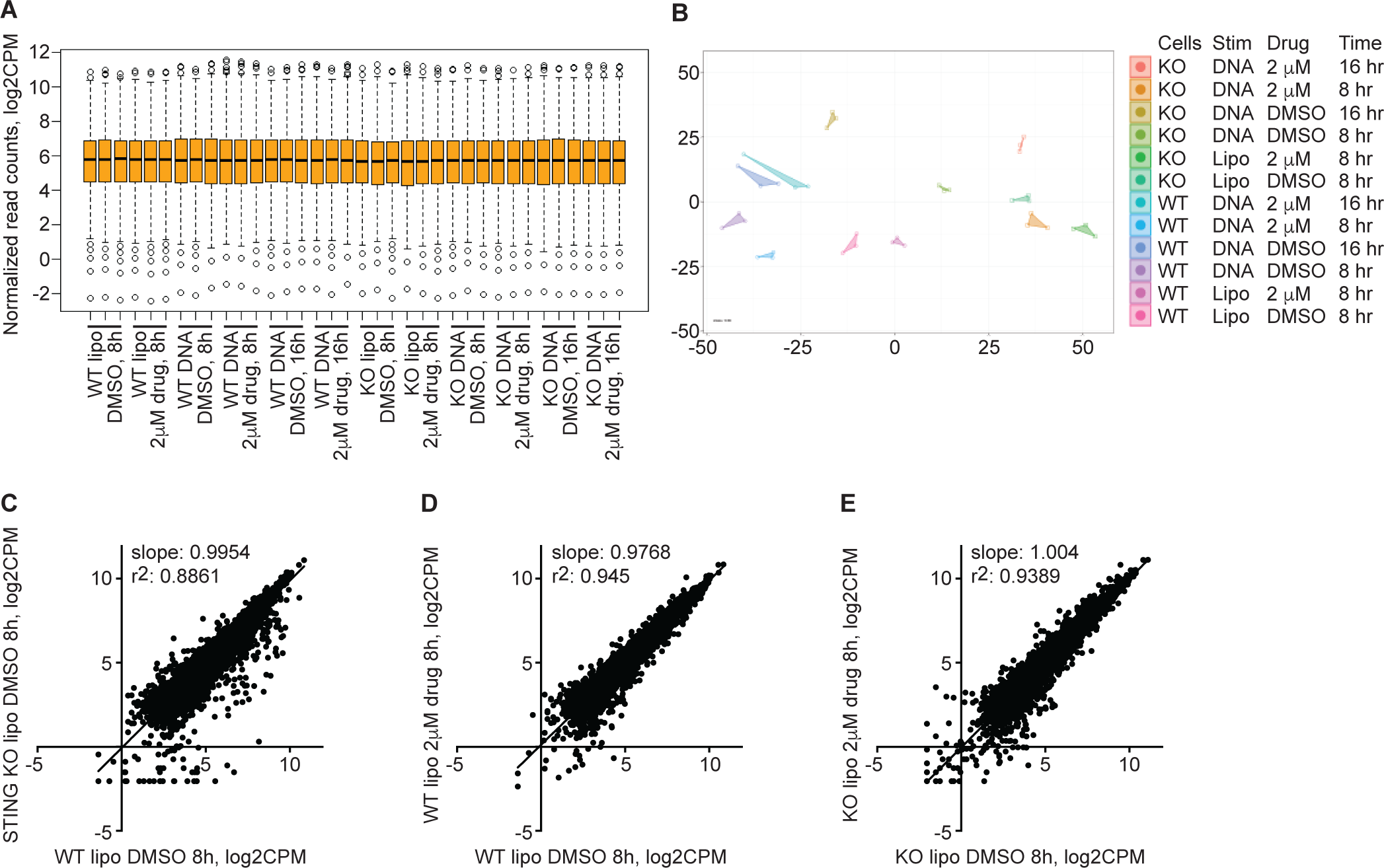
Normalized mRNA-Seq data comparing WT and STING KO U937 cells. (**A**) Boxplot depicting normalized read counts in log2CPM format across all 36 samples. (**B**) A Metrics Dimensional Scaling (MDS) plot, color coded for the three biological replicates of each condition. (**C**) Normalized read counts in log2CPM format, comparing Lipo-treated WT U937 cells to Lipo-treated STING KO U937 cells. (**D**) Normalized read counts in log2CPM format, comparing Lipo-treated WT U937 cells pretreated with DMSO or 2μM N-7441. (**E**) Normalized read counts in log2CPM format, comparing Lipo-treated STING KO cells pretreated with DMSO or 2μM N-7441.

**Figure S5:**
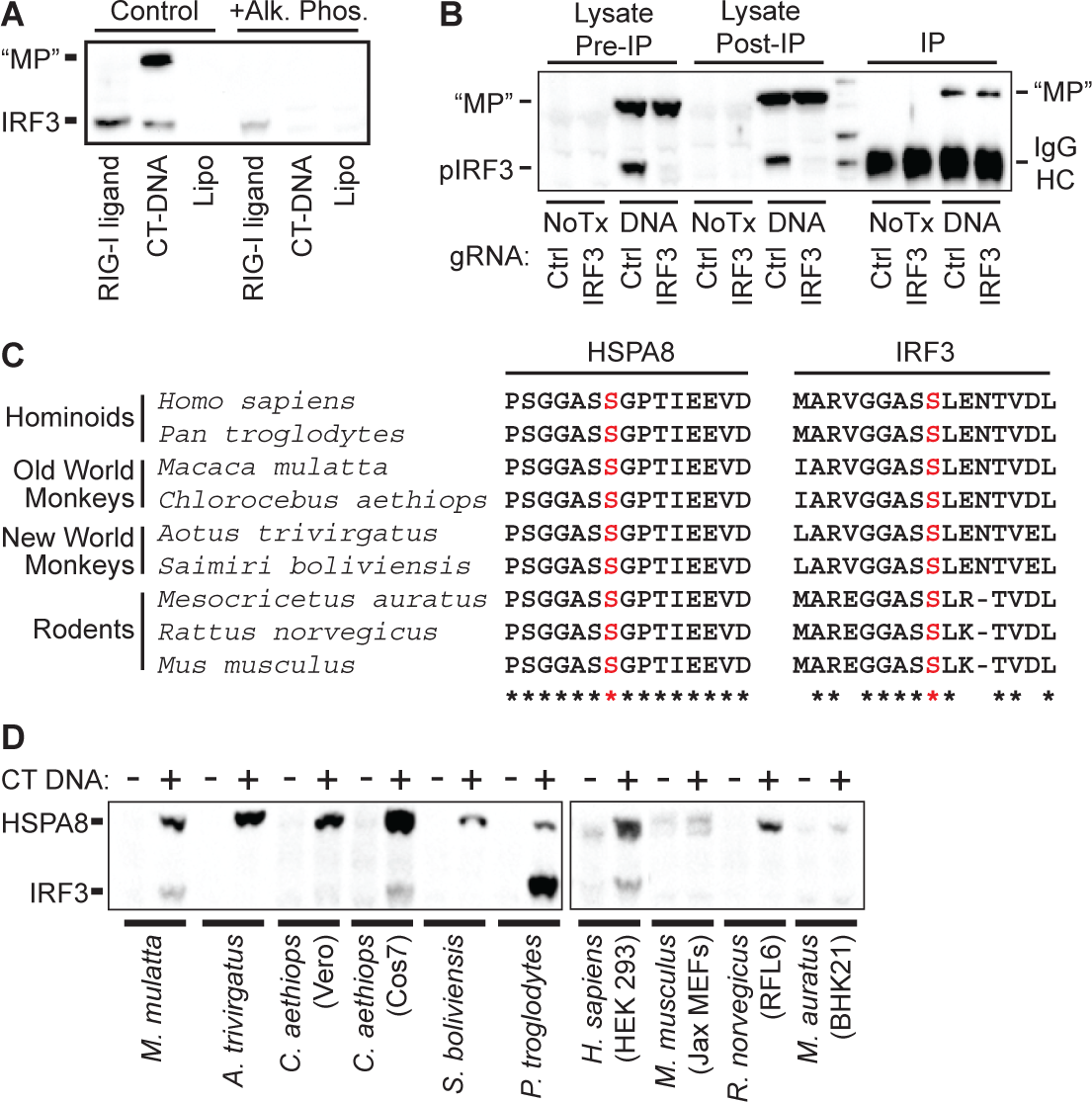
Characterizatin of HSPA8 phosphorylation on serine 638. (**A**) HEK 293 cells were stimulated with CT DNA or RIG-I ligand for 3 hours followed by preparation of extracts that were either left untreated or treated with alkaline phosphatase prior to western blot analysis of IRF3 pS386 and MP. (**B**) Control and IRF3-CRISPR cells were transfected for three hours with CT-DNA, followed by preparation of cell lysates and immunoprecipitation using IRF3 pS386 antibody. Lysates before and after IP and the IP’d material were analyzed by western blot. (**C**) Alignments of HSPA8 and IRF3 amino acid sequences surrounding the phosphorylated serines. (**D**) Cell lines from the indicated species were stimulated with CT DNA for 3 hours, followed by analysis of IRF3 pS386 and HSPA8 pS638 by western blot.

